# Genome assembly and population genomic analysis of *Aegilops caudata* uncover a rich source for disease and insect pest resistance for wheat improvement

**DOI:** 10.64898/2025.12.19.695602

**Authors:** Hong-Lian Ye, Prakitchai Chotewutmontri, Naxin Huo, Han-Chang Chang, Daryl L. Klindworth, Kirk M. Anderson, Sam Gale, Genqiao Li, Fazal Manan, Qijun Zhang, Eric Yao, Swarupa Nanda Mandal, Jennifer Bragg, Amanda Peters Haugrud, Pablo Olivera Firpo, Yue Jin, Xiangyang Xu, Timothy L. Friesen, Zhaohui Liu, Shengming Yang, Shaobin Zhong, Upinder Gill, Subhashree Subramanyam, Marion O. Harris, Taner Z. Sen, Matthew J. Milner, Jan Dvorak, Xiaofei Zhang, Ming-Cheng Luo, Steven S. Xu, Yong Q. Gu

## Abstract

*Aegilops caudata* L. is a rich source of resistance genes to major wheat pathogens and pests, yet its complex genome structure and lack of a reference sequence have hindered genetic analysis, gene discovery, and introgression. Here, we report a high-quality genome sequence assembly of *Ae. caudata* accession S740-69, integrated with optical mapping to precisely delineate introgressed segments from structurally altered genomic regions in wheat-*Ae. caudata* translocation lines carrying a novel stem rust resistance gene. Resequencing and phenotyping of 95 diverse accessions, coupled with k-mer-based association mapping, enabled population-level identification of several novel loci for resistance to wheat pests and diseases, such as greenbug and stem rust. Virus-induced gene silencing confirmed *Aecau6C01G127270* (*SrAect1*) as the first functionally characterized stem rust resistance gene from *Ae. caudata*. This new genomic framework provides a foundation for systematic mining of resistance genes in this species and their efficient introgression into wheat.

## Main

Limited genetic diversity in modern wheat cultivars, resulting from domestication bottlenecks and intensive breeding selection, constrains their resilience to evolving biotic and abiotic stresses. Wild wheat relatives represent a critical genetic reservoir harboring diverse beneficial genes/traits essential for wheat improvement and safeguarding global food security^1–3^. Among them, *Aegilops caudata* L. (syn. *Ae. markgrafii* (Greuter) K. Hammer), a diploid species (2n = 2x = 14, CC genome) naturally distributed across the eastern Mediterranean region^4^, represents a rich reservoir of genes for disease and pest resistance^5–7^, abiotic tolerance ^8,9^, and agronomic traits^10^. *Ae. caudata* contributed the C-genome of tetraploids *Ae. cylindrica* Host (2n = 4x = 28, DDCC) and *Ae. triuncialis* (L.) Á. Löve (2n = 4x = 28, UUCC), which are valuable sources of traits for wheat improvement^11^. Notably, more than 80% of *Ae. caudata, Ae. cylindrica,* and *Ae. truncialis* accessions are resistant to race Ug99 of the wheat stem rust pathogen (*Puccinia graminis* f. sp. *Tritici*), compared to <24% of accessions in the diploid *Aegilops* species with D, M and U genomes ^8,9,12^. Our own evaluations further showed that >77% of accessions tested were resistant to leaf rust, powdery mildew, and Hessian fly, and 40% were resistant to greenbug (Supplementary Table 1), underscoring *Ae. caudata* as one of the richest genetic reservoirs of disease and pest resistance among wheat relatives.

Among the world core collection of *Ae. caudata*, accession S740-69 (Fig. 1a) has been recognized as an exceptionally valuable source of disease resistance since 1984^13,14^. The accession was used to develop a set of six wheat-*Ae. caudata* S740-69 disomic addition lines in wheat cultivar ‘Alcedo’, facilitating introgression of resistance loci against diseases into wheat^5,10,15–17^. We reasoned that availability of a high-quality genome for *Ae. caudata* focusing on this accession, would make this valuable genetic resource more accessible to breeders. *Ae. caudata* chromosomes are extensively rearranged relative to the wheat chromosome macrostructure^17,18^. These rearrangements complicate gene introgression into wheat through recombination induced by *Ph1* locus mutants and result in inconsistencies when identifying *Ae. caudata* chromosomes in different studies ^5,10,17–20^. Genome-wide analysis of structural variants (SVs) is therefore essential for fully exploiting *Ae. caudata* genetic resources ^21,22^.

**Fig. 1.**
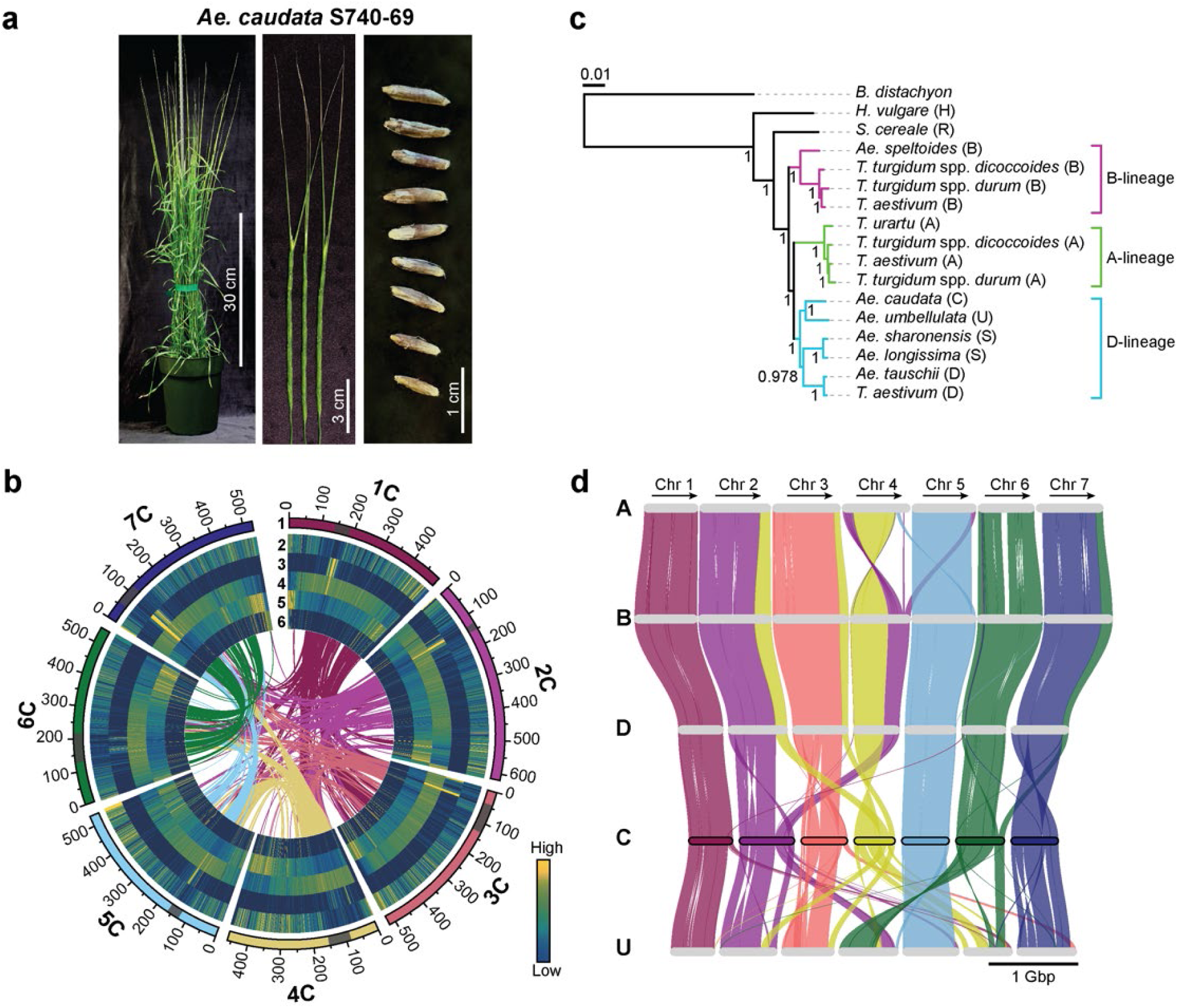
*Aegilops caudata* S740-69 genome assembly and comparison with other Triticeae genomes. **a**, Phenotypic appearance of a whole plant, spikes with the long apical glume awns characteristic of this species, and seeds. **b**, Circos plots showing the genomic features of *Aegilops caudata* with Track 1 showing chromosomes with pericentromeric regions highlighted in grey. Tracks 2 - 6 are heatmap showing the distributions of high-confidence genes (Track 2), Cereba/CRM LTR TEs (Track 3), LTR TEs (Track 4), TIR TEs (Track 5), and resistant gene analogs (Track 6). Relative abundances of each feature per 1 Mbp are plotted in the heatmaps. Connecting lines in the center show inter-chromosome self-syntenic blocks. **c**, Dendrogram depicts the phylogenetic relationships between *Ae. caudata* and other Triticeae species. *Brachypodium distachyon* was used as the outgroup. Genome designations are shown in parentheses. The STAG support values are displayed at the nodes. **d**, Riparian plot showing synteny blocks among the *T. aestivum* (A, B, and D), *Ae. cuadata* (C), and *Ae. umbellulata* (U) genomes. Chromosomes are drawn to scale and display in tandem from left to right. Each chromosome starts from the short arm.

To address these demands, we target *Ae. caudata* accession S740-69 to generate a chromosome-scale genome. We show how a hybrid sequencing-optical mapping approach provides a powerful tool to detect and validate SV in the genome. The initial assembly using PacBio HiFi long reads (∼26x coverage) generated 2,571 contigs with an N50 of 9.2 Mbp. Bionano optical mapping (∼338x) generated 1,350.8 Gbp of optical maps from molecules ≥150 kbp. HiFi-Bionano hybrid scaffolding yielded 184 hybrid scaffolds with an N50 of 43.3 Mbp (Extended Data Fig. 1a). Following gap filling, the draft assembly comprised 176 scaffolds, with an N50 of 44.1 Mbp and a total size of 3.77 Gbp. These scaffolds were then anchored using a linkage map (Supplementary Table 2). The final assembly (Aecau_v1) spanned 3.70 Gbp across 146 anchored scaffolds on the seven chromosomes, with only 0.066 Gbp unanchored (Fig. 1b, Extended Data Fig. 1a-b, and Supplementary Table 3).

We next assessed assembly completeness. k-mer-based method estimated genome size of 3.94 Gbp, while microspectrophotometry reported a larger genome size of 4.7 Gbp^21^. Notably, microspectrophotometry has previously also overestimated *Ae. tauschii* genome by ∼0.8 Gbp^21–23^. Benchmarking Universal Single-Copy Ortholog (BUSCO) analysis against the Poales dataset indicated 98.12% complete orthologs (Extended Data Fig. 1c). These results demonstrate high assembly completeness capturing nearly the entire genome. Assembly continuity was assessed using the LTR Assembly Index (LAI). LAI was 25.62, exceeding the gold-quality cutoff of 20^24^ and placing Aecau_v1 assembly within the top 4% of plant reference genomes^25^. Additionally, seven pericentromeres and fourteen telomeres were detected (Extended Data Fig. 1b). These evaluations establish Aecau_v1 as a nearly complete, high-quality, chromosome-scale reference genome assembly.

For annotation, we first annotated transposable elements which constituted 84.1% of the genome, comparable to other Triticeae genomes (Fig. 1b and Extended Data Fig. 2a). To achieve high-quality gene annotation, we generated full-length transcripts using Iso-seq and RNA-seq from 16 tissues, detecting >120,000 transcript isoforms (Extended Data Fig. 2b). Integrating transcriptomic data with ab initio and protein homology predictions identified 96,320 protein-coding gene models including 43,822 high-confidence (HC) and 43,203 low-confidence genes (Fig. 1b, Extended Data Fig. 2c, and Supplementary Table 4). Gene functions and non-coding genes were annotated, and 1,782 resistance gene analogs (RGAs) were identified (Fig. 1b, Extended Data Fig. 2d-e, Supplementary Table 5-6).

We explored the evolution of *Ae. caudata* by constructing a rooted phylogenetic tree based on HC proteomes. The tree revealed three primary clades corresponding to the A, B, and D lineages (Fig. 1c). The topology of our tree was identical to the tree constructed in the publication of *Ae. umbellulata* assembly, except for the lack of *Ae. caudata* genome at that time^3^. Our tree placed the *Ae. caudata* genome into the D lineage along with the genomes of *Ae. umbellulata, Ae. sharonensis, Ae. longissima,* and the *Ae. tauschii-T. aestivum* D genome pair. While the previous study showed that the U genomes diverged from the D-S common ancestor prior to the D and S split^3^, placement of the new C genome indicates that the C and U genomes shared a common ancestor that diverged from D-S ancestor before their subsequent divergence.

*Ae. caudata* and *Ae. umbellulata* are notorious for their rearranged chromosomes^16–18,26,27^. As the *Ae. tauschii* chromosomes conserved the macrostructure of those of the hypothetical progenitor of the tribe Triticeae^28,29^, its genome is a logical reference in analyses of chromosome structure in the *Triticum-Aegilops* alliance. *Ae. caudata* pseudomolecules 1C and 5C largely conserved their macrostructure. The remaining five pseudomolecules all showed translocation and pericentric inversion events as compared to those of *Ae. tauschii* (Fig 1d, Extended Data 3a). These translocations displaced distally the centromeres of 3C, 4C, and 7C chromosomes (Fig 1b) differing from their Triticeae progenitor chromosomes being metacentric or submetacentric. This finding is consistent with karyotype analyses of *Ae. caudata* genome^18^.

With the Aecau_v1 genome availability, we recently developed codominant markers for the homoeologous group-7 synteny blocks and successfully mapped the stem rust resistance segments in 7AL/6CL and 7DL/6CL translocations^37,38^ in the introgression lines derived from accession S740-69. The identification of introgressed segments using combinational approaches of cytogenetic methods and DNA markers is time-consuming and labor-intensive. To develop reliable and efficient methods, we generated optical maps for four introgression lines. The high-resolution optical maps quickly delineated the boundaries of the introgressed segments at near kilobase-level by aligning them onto the Chinese Spring (IWGSC RefSeq v2.1) and *Ae. caudata* genome assembly (Fig. 2), confirming previous DNA marker mapping results and successful introgressions of *Ae. caudata* 6CL segments into collinear regions in wheat^30^. In addition, optical maps detected other unexpected SVs (Fig. 2a) that had not been detected previously with other methods^30^. Using PacBio HiFi sequencing, we confirmed all the SVs detected in line 6-0015 optical maps and verified the chromosome 6CL terminus at 546.43 Mbp (Fig. 2a), Therefore, Bionano optical mapping is a powerful tool for detailed characterization of alien introgressed lines and their deployments in wheat genetic improvement.

**Fig. 2.**
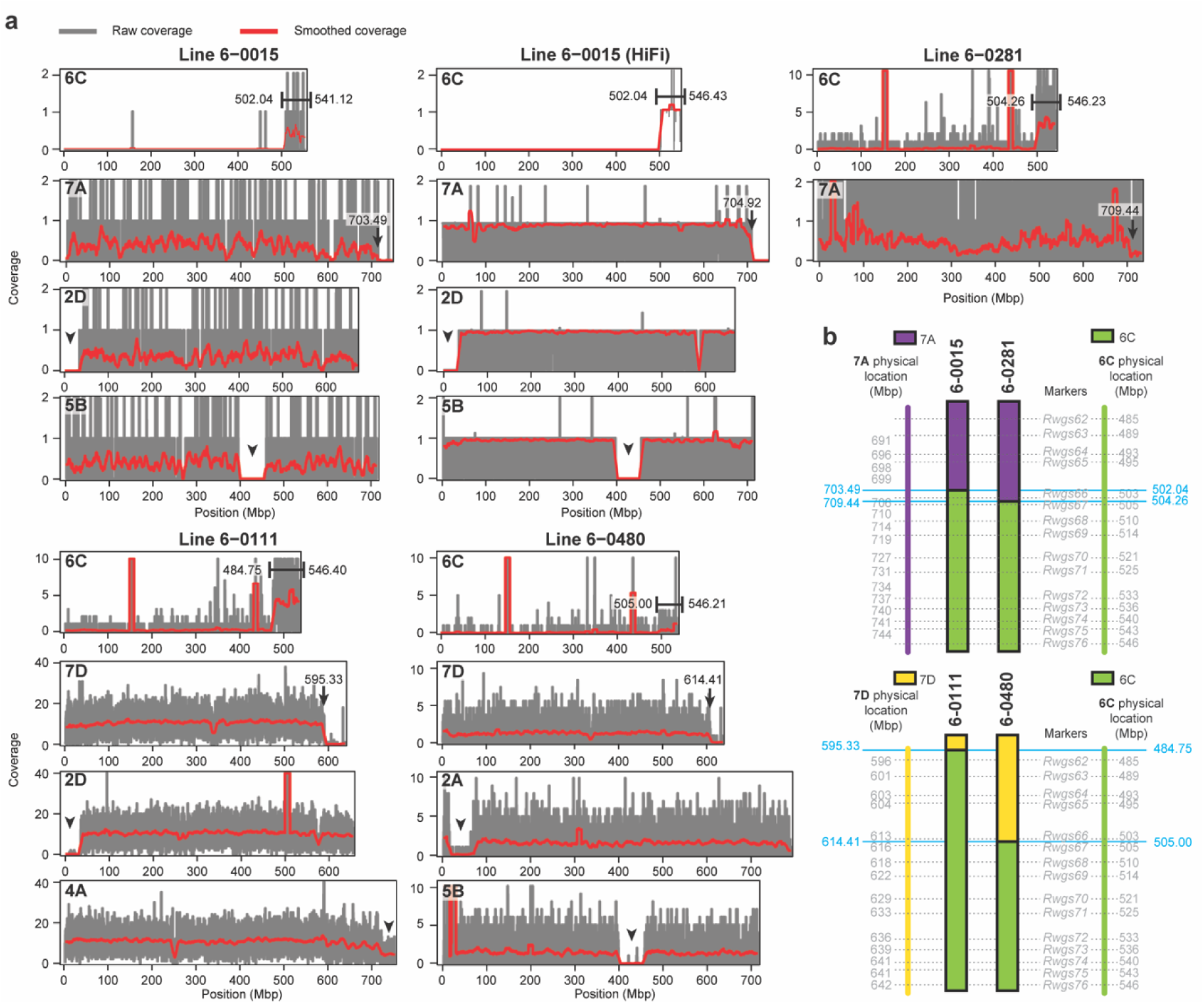
Bionano optical mapping for identification of alien segments and structural variations in the wheat-*Ae. caudata* introgression lines. Four wheat-*Ae. caudata* introgression lines carrying 7A/6C and 7D/6C translocations were analyzed by aligning their optical maps onto the wheat Chinese Spring (IWGSC RefSeq v2.1) and *Ae. caudata* genome assembly **a,** Alignment coverage of optical molecules against Chinese Spring and *Ae. caudata* genomes. Alien segments were detected on chromosome 6C long arm ends, coinciding with the absence of synteny blocks on wheat group-7 chromosome long arm ends. Lines 6-0015 and 6-0281 showed 7A/6C translocations, while 7D/6C translocations were detected in lines 6-0480 and 6-0111. The boundary positions of the segments are marked with bars for chromosome 6C and arrows for wheat chromosomes 7A and 7D. For example, line 6-0015 exhibited a 41.00-Mbp deletion of the wheat 7AL terminus starting at 703.49 Mbp, which was replaced by a 44.39 Mb *Ae. caudata* 6CL terminal segment, starting at 502.04 Mbp on the 6C pseudomolecule. Line 6-0480 exhibited a 28.51-Mbp terminal deletion of 7DL starting at 614.41 Mbp, which was replaced by a 41.43 Mbp-6CL terminus starting at 505.00 Mbp on the 6C pseudomolecule. Some sporadic alignments occurred in the regions with high similarity between Chinese Spring and *Ae. caudata* and these alignments were surrounded by >1 Mbp gaps. Additional structural variations were detected on wheat chromosomes 2, 4, and 5 and marked with arrow heads. PacBio HiFi assembly of line 6-0015 (labelled as HiFi) verified the optical mapping result. **b**, Diagrams of the translocation regions. Chromosomal segments of 6C, 7A, and 7D are shown with the physical positions of previously reported DNA markers. Boundaries of the translocation regions defined by optical genome mapping are highlighted with blue horizontal lines and blue labels. Physical coordinates are given in Mbp.

To study and harness the genetic diversity of *Ae. caudata* and its value for improving modern wheat, we collected a diversity panel of 95 *Ae. caudata* accessions that originated from Greece, Türkiye, Iraq, and of unknown origin (Fig. 3a, Supplementary Table 1), broadly covering the primary distribution range of *Ae. caudata* centered around the Aegean Sea^4^. Whole-genome sequencing (WGS) data was generated, and SNP matrix analysis identified 89 non-redundant accessions with minor residual heterogeneity (Extended Data Fig. 3a-b, and Supplementary Table 7-8). Principal component analysis and population ancestry analysis suggested ∼6 genetic clusters (Extended Data Fig. 3c-e, and Supplementary Table 9). Ancestry coefficients at K=6 aligned well with the geographic origin of the accessions (Fig. 3a). Our reference genome accession AC001 (S740-69) grouped with four other accessions of unknown origin in a distinct genetic clade, preventing identification of its geographic provenance. Two genetic groups were located in Greece, whereas the remaining three were located in Türkiye and Iraq. Ancestry analysis of these accessions at multiple K values (K=4 to K=13) showed mixed ancestries or defined two new genetic groups at the high K values (Extended Data Fig. 3e). This result is consistent with the spike morphological differences observed among different clades (Fig. 3a). Cytogenetic and WGS data analyses revealed that AC007, AC088, and AC094 are tetraploid *Ae. cylindrica* (CCDD), AC044 is *Ae. triuncialis* (CCUU), and AC079 is diploid carried M genome but lacked acrocentric chromosome found in *Ae. caudata* (Extended Data Figure 4a-c).

**Fig. 3.**
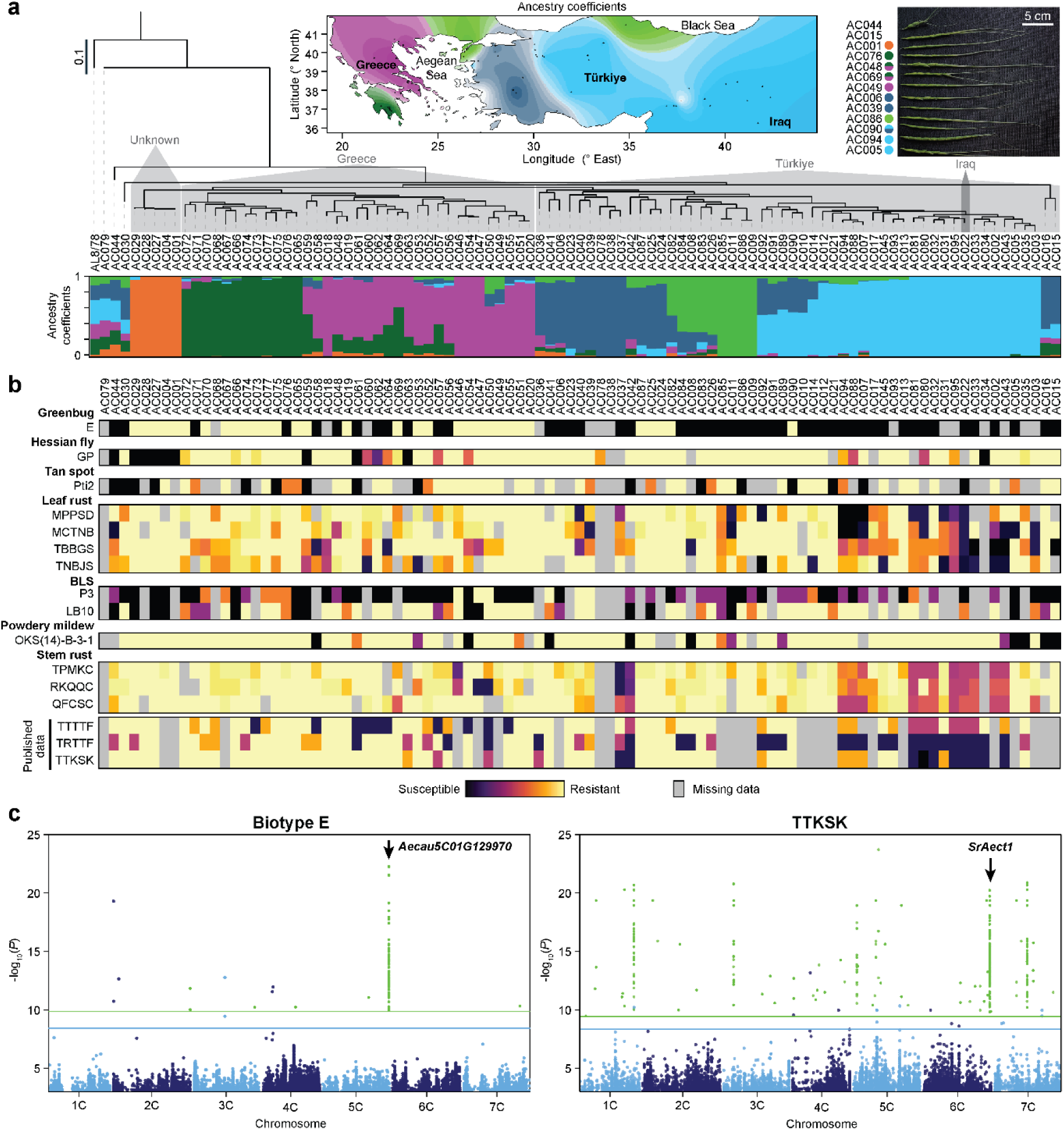
Population structure and disease resistance analysis of *Ae. caudata*. **a**, Population structure analysis. Ancestry coefficients are shown in the bottom heatmap and grouped based on the maximum likelihood tree. Geographic map shows the major ancestry coefficient and the origin of each accession (dot). Images of the spikes from representative accessions for each major ancestry group are shown to the right. Ancestry analysis with other K values is shown in Extended Data Fig. 4e. **b**, Heatmaps of disease resistance phenotypes. Resistance was evaluated for greenbug (*Schizaphis graminum* biotype E), Hessian fly (*Mayetiola destructor* biotype GP), tan spot (*Pyrenophora tritici-repentis* race Pti2), leaf rust (*Puccinia triticina* races MPPSD, MCTNB, TBBGS, and TNBJS), bacterial leaf streak (*Xanthomonas translucens* pv. *undulosa* isolates P3 and LB10), powdery mildew (*Blumeria graminis* f. sp. *tritici* isolate OKS(14)-B-3-1), and stem rust (*Puccinia graminis* f. sp. *tritici* races TPMKC, RKQQC, and QFCSC). Published stem rust phenotypes for races TTTTF, TRTTF, and TTKSK (Ug99) are also included. Two accessions (AC018 and AC039) with an inconsistent greenbug phenotype across replicates is marked as missing data, with raw phenotypic data provided in Supplementary Table 1. Missing values are shown in grey. **c**, k-mer-based GWAS utilized >2 billion k-mers. The significant k-mers are shown with green dots. A major peak on chromosome 5C was detected in greenbug biotype E analysis. GWAS analysis with TTKSK identified six major peaks including the chromosome 6C region within the *SrAect1* locus. SNP GWAS analysis (blue dots) was included for visualization. Horizontal lines show the thresholds of the 5% family-wise error rate (green line for k-mer and blue line for SNP).

We evaluated the *Ae. caudata* diversity panel for resistances against greenbug Hessian fly, tan spot, leaf rust, bacterial leaf streak, powdery mildew, and stem rust. This screening showed that the panel was a rich source of resistance to major wheat pathogens and pests (Fig. 3b, and Supplementary Table 1). For greenbug biotype E, 40% were resistant. A k-mer-based GWAS analysis revealed that 99% of significant k-mers mapped to an NLR gene (*Aecau5C01G129970*) on chromosome 5C (Fig. 3c), in line with greenbug resistance of the chromosome 5C disomic addition line (Supplementary Table 1). This represented the first genetic characterization and mapping of a greenbug resistance gene in *Ae. caudata*.

Screening for reactions to races TPMKC, RKQQC, and QFCSC of stem rust pathogens showed that 95% of accessions were resistant (Fig. 3b) and identified a peak for race RKQQC (Extended Data Fig. 5e). Reactions to races TTTTF, TRTTF, and TTKSK (Ug99) was previously reported^7^. GWAS analysis of TTKSK mapped 5,180 significant k-mers with peaks on chromosomes 1C, 3C, 5C, 6C, and 7C (Fig. 3c). This consists with Alcedo 6C or 7C additional lines exhibiting resistance to TTKSK^31^. The major peak on 6C was mapped to *SrAect1,* a novel resistance locus^30^. Multiple lines of evidence supported *Aecau6C01G127270* as the top candidate within *SrAect1* locus (Extended Data Fig. 6). First, the largest number of k-mers (379) mapped to this gene. Second, the WGS read coverage suggested that the susceptibility was associated with the absence of a ∼2.2 Mbp region containing ∼29 HC genes. Third, RNA-seq analysis across 16 tissue types revealed that *Aecau6C01G127270* was the only one of 29 genes expressed in tissues expected to react to stem rust infection (leaf, leaf sheath, stem). Moreover, *Aecau6C01G127270* encodes a coiled-coil, nucleotide-binding site, leucine-rich repeat protein characteristic of plant resistance genes (Fig. 4a). Virus-induced gene silencing (VIGS) was used to silence the gene expression in two resistant introgression lines. Each of the two constructs targeting *Aecau6C01G127270* reduced the expression by ∼50% (Fig. 4a-b). In control experiments with no virus, empty virus, and non-target gene silencing, all plants remained resistant to TPMKC while *Aecau6C01G127270* silencing plants showed susceptible phenotypes (Fig. 4c). This result confirmed that *Aecau6C01G127270* is equivalent to stem rust resistance gene *SrAect1*. The SRAect1 protein is novel, forming its own clade and grouped with SR45, SR9B, and SR21 proteins (Fig. 4d). Synteny analysis of the *SrAect1* locus showed that the NLR genes, including *SrAect1,* were absent in rice, sorghum, and maize (Fig.4e), suggesting that these genes arose after the divergence of Pooideae from Oryzoideae and Panicoideae, 35-50 million years ago^32^. Notably, the NLR gene was absent from the *SrAect1* locus in *Ae. tauschii* and *Ae. sharonensis,* indicating the highly dynamic nature of this ancient locus.

**Fig. 4.**
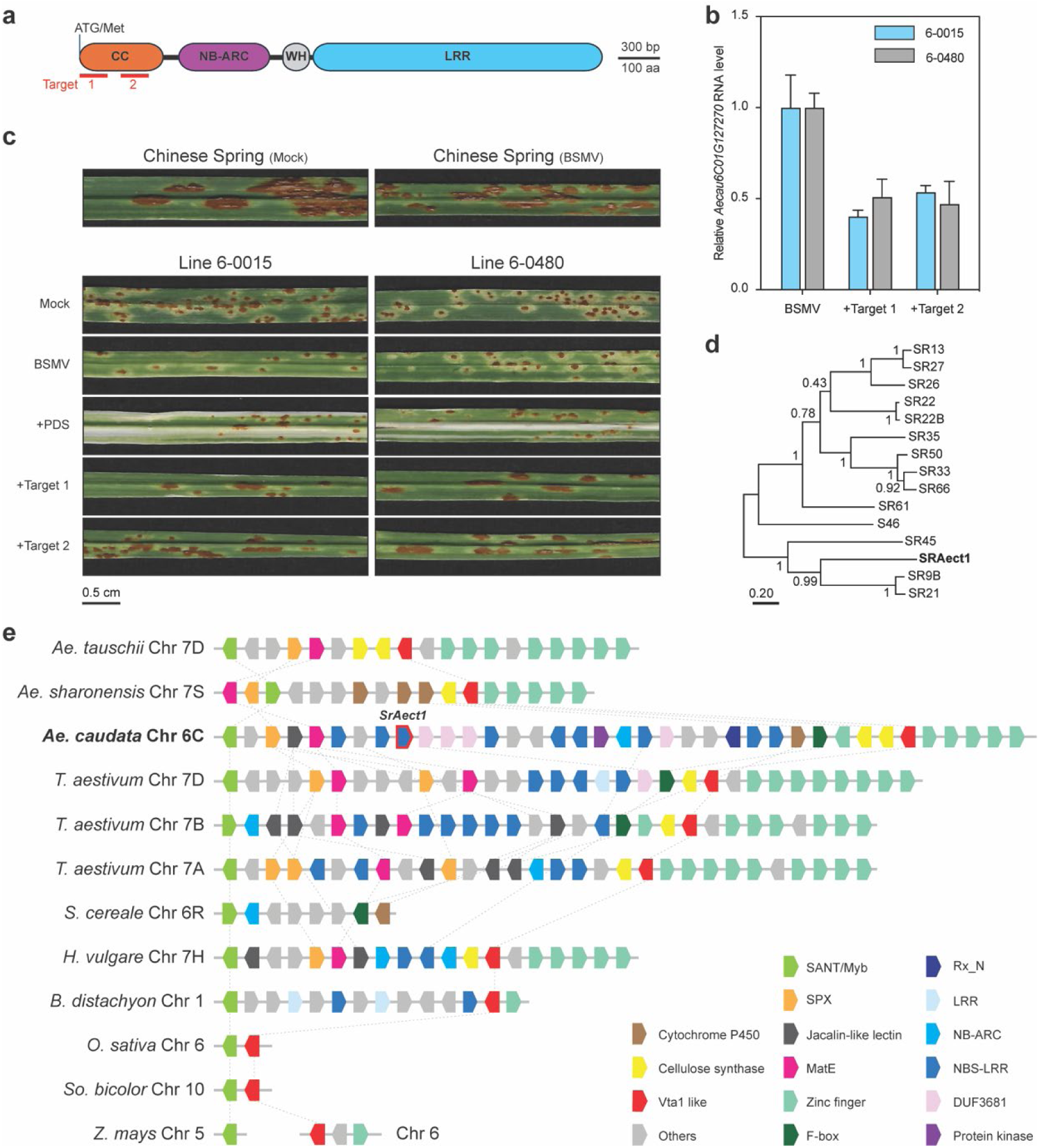
Functional analysis of *SrAect1* (*Aecau6C01G127270*) gene. **a**, Intronless *Aecau6C01G127270* gene encodes a Coiled-coil (CC), Nucleotide-binding site (NB-ARC), Leucine-rich repeat (LRR) protein. Domain structure is drawn to scale. Red bars show the regions of silencing target sequences (Target 1 and 2). WH, winged helix. **b**, RT-qPCR shows the relative level of *Aecau6C01G127270* RNA in the leaves of BSMV-VIGS silencing plants (+Target 1 and 2) compared to empty virus control (BSMV). Means ± SD are shown from 3 biological replicates. **c**, Images showing the representative stem rust TPMKC phenotypes of the VIGS experiment in the two stem rust resistant introgression lines 6-0015 and 6-0480. Chinese Spring was included for comparison. Controls include no virus (Mock), empty virus (BSMV), and non-target gene silencing (+PDS). The *Aecau6C01G127270* silencing plants (+Target 1 and 2) exhibited susceptible phenotypes. **d**, Phylogenetic analysis of SRAect1 protein and the other cloned stem rust resistant (SR) proteins of CC-NBS-LRR type. Bootstrap values are shown at nodes. The SRAect1 grouped with SR45, SR9B, and SR21 proteins. This group is characterized by longer LRR regions. **e**, Synteny analysis of the *SrAect1* loci in grasses. Each box represents a HC gene. Box color indicates protein domain. The *SrAect1* loci are flanked by a gene encoding SANT/Myb domain and a gene encoding Vta1-like domain. Orthologous analysis identified the gene orthologs which are indicated by the vertical dotted lines but was unable to identify *SrAect1* ortholog among the NBS-LRR/NLR genes in the regions. The letters *Ae*, *T*, *S*, *H*, *B*, *O*, *So*, and *Z* in the species names represent genera *Aegilops*, *Triticum*, *Secale*, *Hordeum*, *Brachypodium*, *Oryza*, *Sorghum*, and *Zea*, respectively.

In summary, this study establishes an integrated framework for harnessing genetic diversity from wild relatives. We developed valuable resources including the high-quality assembly of the *Ae. caudata* genome, optical mapping for robust characterization of alien DNA introgression lines, resequencing, population structure, and phenotyping of the collected diversity panel for pest and disease resistance, and functional validation of introgression lines carrying resistance loci. Together, these resources, coupled with k-mer-based association mapping, pave the path from trait discovery to breeding applications and, offer a scalable strategy for accelerating the deployment of valuable traits from untapped wild relatives into elite wheat germplasm, with direct implications for wheat improvement and global food security.

## Materials and methods

### Plant materials

Seeds of *Ae. caudata* accession S740-69 (AC001) used for C genome assembly were provided by Dr. Richard R.-C. Wang at the USDA-ARS, Forage & Range Research Laboratory (Logan, UT, USA). Original seeds for the other 94 *Ae. caudata* accessions in the *Ae. caudata* diversity panel (Supplementary Table 1) were obtained from John Raupp at the Wheat Genetics Resource Center, Kansas State University (Manhattan, KS, USA), and Dr. Harold E. Bockelman at the USDA-ARS National Small Grains Collection (Aberdeen, ID, USA). For seed increase, a single plant from each accession was grown in potting mix under a 16/8 h (light/dark) photoperiod at a temperature setting of 22°C in the greenhouse (Albany, CA, USA). Four weeks after planting, the plants were vernalized at 4°C for eight weeks, then returned to the greenhouse and grown to maturity. Four wheat–*Ae. caudata* S740-69 introgression lines (6-0015, 6-0281, 6-0111, and 6-0480) carrying the stem rust resistance gene *SrAect1* derived from S740-69 were used for Bionano optical mapping of *Ae. caudata SrAect1* introgressions and for *SrAect1* cloning and validation. Lines 6-0015 and 6-0281 carry *SrAect1* within a short 6CL segment in a pair of 7AS·7AL-6CL translocation chromosomes, whereas 6-0111 and 6-0480 carry *SrAect1* on 7DS·7DL-6CL translocations^30^.

### Genome sequencing and assembly

#### DNA sequencing

A single self-pollinated progeny plant of *Ae. caudata* accession S740-69 was grown in potting mix under a 16/8 h (light/dark) photoperiod at 21°C in a growth chamber (Conviron PGR15). Young leaves were collected after a 3-day dark treatment to reduce starch content, immediately frozen in liquid nitrogen, and stored at –80°C until use. Two grams of the young leaf tissue were used to extract high-molecular-weight (HMW) genomic DNA using a protocol optimized for long-read sequencing^33^. DNA quality control was performed using the Agilent FemtoPulse system (Genomic DNA 165kb Kit) for high molecular weight DNA assessment and Agilent 2100 Bioanalyzer (Agilent Technologies) with DNA 12000 Kit for sheared DNA fragment analysis. SMRTbell libraries were constructed using the SMRTbell Express Template Prep Kit 2.0 (PN 101-853-100 version 05, Pacific Biosciences) according to manufacturer specifications. The genomic DNA (5ug) was sheared using the Megaruptor 2 system (Diagenode) generating fragments with an average length of 15-20 kbp. Size-selected libraries were sequenced by Vincent J. Coates genomics sequencing lab at the University of California, Berkeley, on the PacBio Sequel II system using Circular Consensus Sequencing (CCS) mode generating ∼105 Gbp of HiFi reads (∼26X genome coverage).

#### Bionano optical maps

HMW DNA was extracted from young leaves using the Bionano Prep Plant DNA Isolation Kit (Bionano Genomics). DNA was labeled with DLE-1 enzyme and subsequently stained following the DLS G2 protocol (CG-30553-1). Optical maps were generated on the Saphyr genome mapping system using G3.3 chips producing ∼338x genome coverage. De novo assembly was performed using Bionano Solve v3.8.1.

#### Genome assembly

First, PacBio HiFi reads were assembled using Hifiasm (v 0.16.1-r375; parameters: -l 0 -f 39) producing 2,571 contigs^34^. Next, de novo assembly of Bionano reads followed by hybrid scaffolding with the PacBio HiFi contigs were performed using Bionano Solve (v3.8.1) resulting in 184 hybrid scaffolds (3.75 Gbp). Assembly statistics and anchoring process information are provided in Extended Data Fig. 1a and Supplementary Table 3, respectively.

#### Linkage map construction

To enable chromosome-scale genome assembly, we generated a mapping population of 227 F₂ individuals by crossing S740-69 (as the male parent) with a genetically diverse *Ae. caudata* accession, TA2085 (AC006). Genomic DNA was extracted from young leaf tissue at the three-leaf stage of the F_2_ individuals and both parental lines using a modified CTAB method^35^. Genotyping was performed using the Illumina Wheat 90K iSelect SNP assay following manufacturer’s protocols^36^. In total, the assay produced data for 81,587 functional SNPs. SNP calling and clustering were performed using GenomeStudio2.0^37^ (Illumina Inc.) with quality control parameters: call rate ≥ 0.75 and cluster separation ≥ 0.3. This 90K SNP array was designed initially for hexaploid wheat. To ensure accurate allele calling when applied to *Ae. caudata*, manual inspection and adjustment of SNP clustering patterns were performed^38^. Of the total functional SNPs assayed, 6,553 SNPs with indistinguishable clustering patterns were excluded, 59,487 were monomorphic, and 10,712 failed genotypes calling. A total of 4,835 polymorphic SNPs were retained for linkage analysis.

For linkage map construction, manual quality control (QC) was performed in two steps. First, SNPs with >20% missing data across individuals and those showing genotype inconsistencies between parents and progeny were excluded. Second, non-segregating markers were consolidated by retaining one representative marker per group. After these QC and consolidation steps, 708 high-quality SNPs from 222 lines were used for linkage analysis (reduced from the original 4,834 SNPs and 227 lines). Linkage analysis was conducted using QTL IciMapping (v4.2) configured for F_2_ population analysis^39^. Recombination frequencies were converted to genetic distances in centimorgans using the *Kosambi* mapping function, with interval-based marker positioning. SNP markers were grouped using a LOD threshold of 3.0. Map ordering employed K-Optimality and 2-OptMAP algorithms with NN initials of 10. The 90K consensus map served as reference for marker ordering and chromosome assignment. Map refinement was conducted using rippling with a window size of 5. The final linkage map comprised seven linkage groups, each corresponding to one chromosome of the *Ae. caudata* C genome (Supplementary Table 2).

#### Anchoring and gap filling

The 90K SNP linkage map was then used to anchor hybrid scaffolds. The positions of the SNP probes on the scaffolds were determined using Blast+ (v2.15.0; parameter: -word_size 8). Based on the linkage map, 79 scaffolds (3.02 Gbp) were ordered and oriented into draft pseudomolecules. Whole genome alignment with *Ae. tauschii* Aet v5 assembly using minimap (v2 2.24; parameter: -cx asm5) was used to anchor additional 75 scaffolds (0.67 Gbp)^40^. Lastly, the gaps between scaffolds were filled using (parameter: --mode gapfiller) program supplied with the original HiFi read^41^. The final pseudomolecule assembly named Aecau_v1 was composed of 146 anchored scaffolds (3.70 Gbp) in 7 chromosomes and 30 unanchored scaffolds (0.066 Gbp) (Extended Data Fig. 1b).

#### Genome assembly assessment

Jellyfish (v2.3.0)^42^ was used to produce a 21-mer histogram (parameters”count -C -m 21” followed by “histo -h 3000000”). Genome size estimation based on the histogram was performed using findGSE^43^, a k-mer based method in R (v4.5.0). Assembly completeness was evaluated using BUSCO (v5.4.5) with the Poales dataset (Poales_odb10) and default settings^44^. To assess assembly continuity, the LTR Assembly Index (LAI) was calculated using LTR_retriever (v3.0.4; parameter: - q), based on the intact LTR elements identified by EDTA during TE annotatio^24^ described below. Pericentromeric regions were delineated from *Ae. tauschii* Aet v4 coordinates^29^ by whole genome alignment^40^. Telomeric regions were identified using the TIDK toolkit^45^ with parameters “search -s AAACCCT” and with three counting window sizes (parameter: -w 0.01, 0.1, and 1 Mbp). Final telomeric sites were manually defined from TIDK outputs as regions with telomeric repeat counts exceeding 30-fold the genome-wide average within the 0.01 Mb window.

### Gene annotation

Annotation was performed using a modified procedure derived from the previously published methods for BRAKER pipeline^46^ and the recent einkorn assembly^47^.

#### Transcriptome sequencing

To assist annotation, Iso-seq and RNA-seq data were generated from 16 tissue types of S740-69 (Extended Data Fig. 2b). Total RNA was extracted from seedling leaf, seedling crown, stem before boot, leaf sheath, glume after heading, flag leaf, rachis, seedling root, young spike, spike after heading, peduncle, anthers after heading, spikelets, seeds 14 days post anthesis, seeds 20 days post anthesis, and leaf after 2 months of cold treatment. In brief, ∼100 mg of ground sample powder from each tissue was used for RNA isolation using the RNeasy Plant Mini Kit (Qiagen) according to the manufacturer’s protocol. RNA quality and Integrity were assessed using Agilent Bioanalyzer. RNA-seq libraries were prepared from each tissue using oligo-dT selection and sequenced using Illumina NovaSeq 6000 in 150 bp paired-end mode by Novogene Corporation Inc., producing a total of over 137 Gbp data. For Iso-seq, RNA samples were pooled into 2 pools. Library preparation (PacBio SMRTbell prep kit 3.0), sequencing (PacBio Sequel II in CCS mode), and data processing (SMRT Link v12.0.0.177059) was performed by the Genome Center, the University of California, Davis, producing >127k transcript isoforms from each pool.

#### Transposable element identification

De novo transposable element annotation was performed using the Extensive de novo TE Annotator (EDTA, v1.9.4; parameters: --sensitive 1 --anno 1 --evaluate 1) with the coding sequences of *Ae. tauschii* (NCBI RefSeq accession GCF_002575655.2; parameter: --cds) used during gene sequence purging (Extended Data Fig. 2b)^48^. Additional masking was performed on the EDTA masked genome using the Tandem Repeat Finder (v4.10.0rc2; parameter: 2 7 7 80 10 50 500 -d -m -h)^49^.

#### Protein-coding gene annotation

Gene annotation was performed by integrating transcriptome data, ab initio predictions, and protein homology evidence. For genome-guided transcriptome assembly, RNA-seq reads were trimmed with cutadapt 4.0^50^ and aligned to the genome using HISAT2 (v2.2.1; parameter: --dta)^51^. The aligned BAM files were used to generate transcript assembly with StringTie^52^ (v2.2.0; parameters: -m 150 - f 0.3 -t) and produced the GTF files. For Iso-seq, the high-quality reads from both RNA pools were aligned to the genome using mimimap2 (v2.24; parameter: -ax splice:hq -uf - -secondary=no)^40^ and the transcript isoforms were collapsed using cDNA_Cupcake (v29.0.0; parameters: collapse_isoforms_by_sam.py --dun-merge-5-shorter) (https://github.com/Magdoll/cDNA_Cupcake). The RNA-seq and Iso-seq transcripts were merged using StringTie (v2.2.0; parameters: --merge -m 150)^52^. De novo transcriptome assembly was performed using Trinity (v2.15.1)^53^ using the RNA-seq and Iso-seq BAM alignment files with parameters: --genome_quided_bam --long_reads_bam --genome_guided_max_intron 10000. PASA (v2.5.3)^54^ was used to annotate the genome with the de novo transcriptome. Coding regions in the transcripts were predicted using TransDecoder (v5.7.1; https://github.com/TransDecoder/TransDecoder) guided by Pfam-A HMM models (Pfamv37.0) and all protein in Swiss-Prot (UniProt release 2025_03).

Ab initio gene prediction was conducted using BRAKER3^55,56^ which utilized AUGUSTUS and GeneMark predictions. The BRAKER3 was run as described previously^46^ on the genome masked by EDTA and TRF, incorporating aligned RNA-seq and Iso-seq BAM files, as well as proteomes from closely related species obtained from NCBI (*T. dicoccoides*, GCF_002162155.2; *T. urartu*, GCF_003073215.2) and from original publications: *Ae. tauschii* (Aet v6.0)^23^, *Ae. umbellulata* (TA1851_v1)^3^, and *T. aestivum* (IWGSC Refseq v2.1)^57^. The GTF outputs from BRAKER3 RNA-seq and Iso-seq predictions were merged using TSEBRA (v1.1.2.5)^58^ supplied with the BRAKERS3 hintsfile files of the proteomic evidence, and parameter “--filter_single_exon_genes” to remove BRAKER3 false positive single exon gene models lacking proteomic evidence.

Protein homology-based gene prediction was performed using miniprot (v0.13-r248; parameters: --gff -I --outc=0.7), incorporating proteomes from NCBI genomes (*Ae. tauschii* GCF_002575655.2, *T. aestivum* GCF_018294505.1, *T. urartu* GCF_003073215.2, *T. dicoccoides* GCF_002162155.2, *B. distachyon* GCF_000005505.3, *H. vulgare* GCF_904849725.1), EnsemblPlants (*T*. *aestivum* ‘ArinaLrFor’ PGSBv2.1), the HC gene set of *T. aestivum* ‘Kariega’^57^, all Triticeae and Poaceae proteins in UniProt (Release 2025_03).

All structural predictions were integrated using EVidenceModeler (v2.1.0) where the weights for transcriptome, MAKER3, and protein homology evidence were set to 12, 2, and 1, respectively^54^. Finally, the gene annotation was refined through two rounds of updating the gene models from the EVidenceModeler output with PASA (v2.5.3) using RNA-seq and Iso-seq data^54^.

Protein-coding gene models were classified into 2 categories, high confidence (HC) and low-confidence (LC), following published protocols ^3,59,60^. Each predicted model was matched against three protein datasets: the TE database, PTREP release 19 (https://trep-db.uzh.ch/), Poales proteins in UniProtKB and the curated Magnoliopsida proteins from Swiss-Prot (UniProt release 2025_03). Search was performed using DIAMOND (v2.1.13)^61^, with an E-value below 1 x 10^-10^. A minimum query coverage of 95% was required for PTREP hits, and a minimum target coverage of 95% and percentage identity of 66% for UniProtKB and Swiss-Prot hits. In addition, all models were evaluated using BUSCO (v5.7.1) with the Poales_odb10 lineage dataset^44^. A model was considered complete if it began with a Methionine (start codon) and ended with a stop codon^62^. HC models were defined as complete protein models that had no hits in the TE database PTREP but showed at least one hit in any of the other datasets (Poales UniProtKB, Magnoliopsida Swiss-Prot, Poales_odb10 BUSCO). LC models were defined either as for incomplete protein models that showed no PTREP hits but had at least one hit in other datasets, or complete protein models that showed no hit in any datasets (Extended Data Fig. 2c).

#### Non-coding gene annotation

These genes were annotated using cmscan function from Infernal (v1.1.4; parameters: --cut_ga --rfam --nohmmonly --oskip --fmt 2) based on the Rfam covariance model release 14.10 (Extended Data Fig. 2d)^63^.

#### Disease resistance gene analog identification

Resistance gene analogs (RGAs) were identified from the HC protein set using RGAugury supplied with Pfam-A HMM models (Pfam37.0)^64^ and NLRtracker^65^. For NLRtracker, HC proteins were first annotated using InterProScan (v5.72-103.0; parameters: -f gff3 -t p -appl Pfam, Gene3D, SUPERFAMILY, PRINTS, SMART, CDD, ProSiteProfiles)^66^. Additional annotation for the supplied NLRtracker NLR motif was performed using FIMO (v5.5.7)^67^. The output files were passed to NLRtracker together with HMMER (v3.3.2) in R (v4.4.1). Identified RGAs are summarized in Supplementary Table 6.

### Genome comparison

#### Phylogenetic analysis

The HC proteome of the C genome was compared to 12 proteomes: two proteomes from Phytozome 14 (*B. distachyon* v3.2 and *H. vulgare* Morex V3 primary transcripts); one HC proteome from GrainGenes (*S. cereale* Rye_Lo7_2018_v1p1p1); two proteomes from NCBI (*T. dicoccoides* GCF_002162155.2 and *T. turgidum* GCA_900231445.1); a proteome from Ensembl Plant release 62 (*T. urartu* IGDB); and six additional proteomes from the original publications (*Ae. umbellulata* TA1851_v1, *Ae. speltoides*, *Ae. sharonensis*, *Ae. longissima*, and *T. aestivum* IWGSC Refseq v2.1) ^3,57,68^. For each gene, only the longest isoform was retained. Species tree was constructed using OrthoFinder (v3.0.1b1) with the “-M msa” parameter^69^.

#### Synteny analysis

Self-synteny analysis was performed using the longest protein isoform from each HC gene. BlastP was performed using DIAMOND (v2.1.13)^61^ with parameters “--evalue 0.00001 --outfmt 6” and the synteny blocks were determined using MCScanX (v1.0.0)^70^ with “-e 0.00001” parameter. To compare the A, B, C, D, and U genomes, the longest protein isoforms from *Ae. umbellulata* (TA1851_v1)^3^, *T. aestivum* (IWGSC Refseq v2.1)^57^, and the C genome was analyzed using GENESPACE (v1.3.1)^71^ in R (v4.4.1). GENESPACE analysis used MCScanX (v1.0.0), OrthoFinder (v2.5.5), and DIAMOND (v2.1.8) for orthology and synteny inference.

#### Other comparisons

Comparison of the TE annotation was conducted using the published results of *T. aestivum* (IWGSC CS RefSeq v2.1)^57^, *Ae. tauschii* (Aet v4.0) ^29^ and *Ae. umbellulata* (PI 554389)^3^. For RGA comparison, the HC protein sequences of these genomes were analyzed with RGAugury^64^ and NLRtracker^65^ as described above.

### Analysis of wheat-*Ae. caudata* introgression lines

#### Bionano optical map

Bionano optical maps were generated as described above for four wheat-*Ae. caudata* introgression lines (6-0015, 6-0281, 6-0111, and 6-0480). De novo assembly was performed for 6-0015 data as described above using Bionano Solve (v3.8.3).

#### PacBio HiFi assembly

Data was generated for 6-0015 as described above using PacBio Revio system generating ∼575 Gbp of HiFi reads (∼37x genome coverage). Assembly was performed using Hifiasm (v0.23.0-r691) with parameters “-l0 -f39” producing 9,222 contigs with N50 of 50.29 kbp and a total length of 14.912 Gbp^34^. Scaffolding was performed with scaffold command (parameters: –mm2-params ‘-x asm5 -f 0.05’) from Ragtag v.2.1.0^72^ using IWGSC RefSeq v2.1^57^ assembly and minimap2 2.4 aligner^40^.

### Detection of introgression of *Ae. caudata* into wheat

The optical map assembly of 6-0015 and the raw optical molecules of the other lines (6-0281, 6-0480, and 6-0111) were aligned to the Chinese Spring (IWGSC RefSeq v2.1) and *Ae. caudata* (Aecau_v1) genomes using Bionano Solve (v3.8.2). Key alignment parameters included a *P*-value thresholds of 1 x 10^-10^ and sample-optimized false positive rates. Bionano coverage depth was calculated by counting the number of independent molecule alignments at each genomic position, providing a direct measure of alignment density for detecting structural variations. The HiFi assembly (6-0015) was aligned to the Chinese Spring (IWGSC RefSeq v2.1) and *Ae. caudata* (Aecau_v1) genomes using minimap2 (v2.26; parameters: -f 0.02 -cx asm5)^40^. The HiFi alignments shorter than 50 kbp or with mapping quality lower than 30 were excluded. To help visualize structural variations, the smoothed coverage lines were generated using a 1,000-site sliding window averaging, and coverage data were visualized using R (v4.3.2) with the ggplot2 and zoo packages.

#### Translocation boundary

The regions with shifts in the smoothed coverage lines were examined manually. In optical alignment to *Ae. caudata* (Aecau_v1), only the alignment regions with confidence scores higher than the chromosome average confidence scores plus three times of standard deviations were considered true alignments. The second criterion is the gap size between alignment sites. The alignment was considered as connected if the gap size is not larger than the chromosome average gap size plus two times of its standard deviations. For HiFi assembly alignment, alignments only longer than 50 kbp with the highest mapping quality at 60 were considered true alignments. The boundary was determined based on the positions of the true alignments with connected coverage.

### *Ae. caudata* diversity panel analysis

#### Sample preparation and whole-genome sequencing

To characterize genome-wide diversity and population structure, we acquired all the available *Ae. caudata* accessions from the two U.S. collections, the Wheat Genetics Resource Center and the National Small Grains Collection, for a total of 95 accessions (Supplementary Table 1). This panel represents natural populations from Iraq, Greece, Türkiye and some accessions with unknown origin. Seeds were planted in cone containers and cold-treated at 4°C for 7 days. After six weeks of growth, approximately 450 mg of fresh leaf tissue was collected and immediately frozen in liquid nitrogen. Genomic DNA was extracted using a modified CTAB protocol with RNase A treatment to remove residual RNA^35^. DNA libraries were prepared and sequenced on the Illumina NovaSeq X Plus using 150-bp paired-end mode for ∼10x genome coverage by Novogene, Inc.

#### SNP calling

The WGS reads were trimmed for quality (“-q 10”) and adapter removal with cutadapt 4.8^50^ and aligned to Aecau_v1 genome using HISAT2 (v2.2.1)^51^. The duplicate reads were removed from the alignment files using samtools 1.17 markdup function^73^. SNP call was performed using bcftools (v1.19) with “mpileup --skip-indels --max-depth 10000 -q 20 -Q 20 -P ILLUMINA -a DP, AD | call -mv -a GQ | filters -g 5 | view --types snps -m 2 -M 2” parameters^74^. SNPs were filtered again using filter_vcf.awk (https://bitbucket.org/ipk_dg_public/vcf_filtering) with parameters “-v dphom=2 -v dphet=4 -v minqual=40 -v mindp=100 -v minhomn=1 -v minhomp=0.9 -v tol=0.2 -v minmaf=0.01 -v minpresent=0.8” producing 46.4 million SNPs for redundancy and population analysis and with parameters “-v dphom=2 -v dphet=4 -v minqual=40 -v mindp=100 -v minhomn=1 -v minhomp=0.3 -v tol=0.2 -v minmaf=0.05 -v minpresent=0.9” for GWAS analysis.

#### Redundancy and residual heterogeneity

Following a previously published method^75^, five million SNPs were randomly sampled from the SNP matrix in five independent replicates. Pairwise SNP identity was calculated for all accession pairs, and those with an average identity exceeding 99.5% were considered redundant (Supplemental Table 8). Based on this criterion, two samples of S740-69 (AC001 and AC004) originating from different sources were classified as redundant (Supplemental Table 1). The final set of 89 non-redundant accessions are described Supplemental Table 1.

Residual heterogeneity was calculated using a previously published script based on the full biallelic SNP matrix (46.4 M)^75^. Except for the two (AC001 and AC004) accessions, all accessions had heterogeneity values well below the 0.1 threshold (Supplementary Table 7). The highest residual heterogeneity observed was 0.07.

#### Population structure and phylogenetic analysis

To assess population structure, we incorporated *Ae. tauschii* accession AL8/78 WGS data (BW_01192)^75^ as an outgroup for comparison alongside our *Ae. caudata* panel. A total of five million SNPs was sampled from the full SNP matrix for downstream analysis. Principal component analysis (PCA) and the admixture coefficient inference were performed using R package LEA (v3.20.0)^76^. The geographical map was generated using tess3r^77^. A maximum likelihood tree was generated busing IQ-TREE 2 (v3.0.1) with “-m GTR+ASC -B 1000 -bnni -t PARS” parameters^78^.

#### Phenotype evaluation

For insect resistance, the *Ae. caudata* was screened for resistance to greenbug (*Schizaphis graminum* biotype E) and Hessian fly (*Mayetiola destructor* biotype GP), following established protocols^79^ with 81 and 87 accessions yielding reliable phenotypic data, respectively. *Ae. caudata* accessions were selected to evaluate their resistance to tan spot (*Pyrenophora tritici-repentis*) (race Pti2) and bacterial leaf streak (*Xanthomonas translucens* pv. *undulosa*) (isolates P3 and LB10) following published protocols^80,81^. The panel was evaluated for resistance to wheat leaf rust using four *Puccinia triticina* races (MPPSD, MCTNB, TBBGS, and TNBJS) following the protocol described previously^82^. For powdery mildew, the *Ae. caudata* panel was assessed for resistance to isolate OKS(14)-B-3-1 of *Blumeria graminis* f. sp. *tritici* at the USDA-ARS Peanut and Small Grains Research Unit (Stillwater, OK, USA) following the protocol described previously^83^. Disease reactions were evaluated 10 days after inoculation and classified into four infection type (IT, 0-4) categories based on symptom development and sporulation levels: highly resistance (HR, IT = 0-1), moderately resistance (MR, IT = 2), moderately susceptible (MS, IT = 3), and highly susceptible (HS, IT = 4).

Stem rust resistance in the *Ae. caudata* diversity panel was assessed using three *Puccinia graminis* f. sp. *tritici* (*Pgt*) races (TPMKC, RKQQC, and QFCSC). Seedlings were inoculated and then incubated under controlled greenhouse conditions (20-23°C, 16/8 h photoperiod) following established protocols^84^. Infection types were evaluated 13-14 days after inoculation using the Stakman^85^ scale where: IT = 0 (immune), IT = 0; (nearly immune), IT = 1 (very resistant), IT = 2 (moderately resistant), IT = 3 (moderately susceptible), and IT = 4 (very susceptible). Phenotypic data for other *Pgt* races (TTKSK, TRTTF, and TTTTF) was obtained from a previously published study^7^. For quantitative genetic analysis, all infection type scores (0-4 scale) were converted to a 0-9 numerical scale using a published python script^86^.

#### Cytogenetics analysis

Root tips (1-2 cm) were collected from germinated seeds and treated with N₂O at 10 atm (150-170 psi) for 2 hours as previously described^87^. Root tips were fixed in ice-cold 90% acetic acid for 10 min and stored in 3:1 fix solution for two days then (ethanol: glacial acetic acid) at -20°C. Genomic in situ hybridization (GISH) and Fluorescence in situ hybridization (FISH) were conducted as described previously^88^. For chromosome counting, metaphase spreads were stained with antifade mounting medium containing propidium iodide (Vector Laboratories, Burlingame, CA) and imaged using an Axiocam 506 mono camera mounted on a Zeiss Axio Imager M2 microscope. Chromosome number was determined from well-spread metaphase cells.

### Comparison of WGS data to other *Aegilops* genomes

To determine subgenome compositions of each accession, the WGS data were aligned to various *Aegilops* genomes using BWA-mem2 (v 2.3)^89^ with parameters “mem -B 10” limiting mismatch rate to 0.001821. The number of exactly matched reads was determined using samtools 1.17 ^73^ with parameters “view -F 2304 -d NM:0” and converted to the percentage of total reads. Reference genomes included: *Ae. caudata* (Aecau_v1; C genome), *Ae. tauschii* AL8/78 (Aet v6; Lineage 2 D genome; GCF_002575655.3), *Ae. tauschii* TA10171 (Lineage 1 D genome)^1^, *Ae. comosa* PI 551049 (M genome; GCA_044584955.1), *Ae. ventricosa* RM271 (N genome was extracted)^90^, *Ae. longissima* AEG-6782-2 (S genome)^68^, *Ae. umbellulata* TA1851 (U genome; GCA_032464435.1). For validation, raw sequencing data from *Ae. tauschii* AL8/78 (NCBI SRA accession SRR13363412) and other tetraploid *Aegilops* accessions were also included: *Ae. cylindrica* AE 656 (ERR7459049 and ERR7459051), *Ae. cylindrica* AE 1594 (SRR27914108), *Ae. juvenalis* AE 497 (SRR27914090), *Ae. ventricosa* AE 897 (SRR27914124), and *Ae.* triuncialis AE 659 (ERR7459062 and ERR7459063) and AE 459 (SRR27914112), *Ae. longissima* AE 334 (SRR27914225), *Ae. peregrina* AE 547 (SRR27914167), *Ae. kotschyi* AE 120 (SRR27914160), and *Ae. biuncialis* AE 1184 (SRR27914101).

#### SNP-based GWAS

To provide genomic context for k-mer GWAS results, a SNP-based association analysis was performed using SNPs called from the same WGS dataset. SNPs were filtered stringently, retaining only those with a minor allele frequency (MAF) ≥0.05, minimum present rate ≥90%, and maximum heterozygosity ≤0.7, resulting in a set of 11,267,846 high-quality SNPs.

GWAS for resistance to greenbug biotype E and stem rust pathogen race TTKSK were performed using five models (GLM, MLMM, FarmCPU, BLINK, and SUPER) implemented in the GAPIT^91^ (v3.0) package in R (v4.3.2). Based on QQ plot assessment (*P* ≤ 0.000001), the BLINK model showed the best fit for both analysis. The SNPs with - log_10_ (*P*) values were used solely to provide a visual reference for the k-mer based Manhattan plots.

#### k-mer-based GWAS

The trimmed WGS reads generated above and the phenotype values were used in the analysis with a published pipeline with default parameters^92^. The significant k-mers were identified with 5% family-wise error rate. We did not detect a major association peak, except for powdery mildew and stem rust resistance (Extended Data Fig. 5).

### Identification and verification of a stem rust resistant candidate gene

#### *SrAect1* gene identification

The significant k-mers associated with stem rust resistance were aligned to the *Ae. caudata* (Aecau_v1) genome using bowtie (v1.3.1; parameters: - -all --strata –best) to determine genomic positions^93^. The WGS read coverage at the gene level was determined from the WGS alignment files using FeatureCounts^62^ to visualize the chromosomal deletion regions. The RNA-seq alignments generated during the annotation were used to determine the tissue specific expression of each gene using FeatureCounts read count. The gene read counts were normalized using the TPM method^94^. This analysis pointed to *Aecau6C01G127270* as a candidate gene for the *SrAect1* locus.

#### Virus-induced gene silencing

Silencing of the target gene in the introgression lines (6-0015 and 6-0480) were performed using the system based on barley stripe mosaic virus (BMSV)^95^. The BSMV system was composed of 3 constructs for each of the tripartite genome (α, β, and γ). The silencing constructs were generated by addition of the target sequence in reverse orientation after the γb stop codon in pCB301-BSMVγ plasmid. As a positive visual control, a conserved fragment of wheat phytoene desaturase (PDS) gene (whole exon 6 of *TraesCS4B03G0788200*; 156 bp) was used. This PDS fragment shows high sequence conservation with the two other PDS homologs, differing by only one nucleotide. To silence the stem rust gene candidate *Aecau6C01G127270*, two non-overlapping target sequences positions 1-200 bp and 300-500 bp from the start codon were used for Target 1 and Target 2, respectively. BLASTn (Blast+ v2.15.0) found no hit to other genes. The recombinant plasmids were transformed into *Agrobacterium tumefaciens* EHA105. Following a previously published method^96^, virus was produced in *Nicothiana benthamiana* and inoculated onto the first leaf of wheat seedlings at the two-leaf stage. The treated plants were maintained in a growth chamber (Conviron PGR15) under cycles of 16-h light at 21 °C/8-h dark at 18 °C cycles.

#### Stem rust inoculation and disease scoring

After 14 days of VIGS treatment, the urediniospores of the *Pgt* race TPMKC were suspended in non-phytotoxic paraffinic oil and sprayed onto the wheat leaves. Inoculated plants were misted with water and covered with a misted transparent plastic bag and incubated in a growth chamber (Conviron PGR15) at 21 °C with darkness for 12 h, followed by 6 h under light. The bag was then removed, and plants were maintained under a 16-h light/8-h dark cycle at 21 °C in the growth chamber. Disease phenotypes were scored 14 days after inoculation using the Stakman^85^ infection type scale. Phenotypic images were captured using a scanner (Color LaserJet Pro MFP 4301fdn, Hewlett-Packard).

#### RT-qPCR

RNA was isolated from leaves 14 days after VIGS treatment using the RNeasy Plant Mini Kit (Qiagen). DNA was removed using the TURBO DNA-free Kit (Thermo Fisher Scientific, San Francisco, CA, USA). RT-qPCR was performed using iScript One-Step RT-PCR Kit with SYBR Green (Bio-Rad, Hercules, CA, USA) on QuantStudio 3 Real-Time PCR System (Thermo Fisher Scientific). The absence of DNA was confirmed in the reaction without reverse transcriptase. The published primers for *CJ705892* gene were used as a reference control^97^. The *Aecau6C01G127270* primers are 5’-TCCCGGTAGTGATGGTCCAA and 5’-CGGACACGGGGACACATAAA. The thermal cycling conditions included RT at 50 °C for 10 min, followed by denaturation at 95 °C for 5 min, and then 40 cycles of 95 °C for 15 sec and 60 °C for 1 min. The single product was validated by melting curve analysis at the end of each RT-qPCR. Two technical replicates of each of the three biological replicates were performed. Amplification efficiencies were determined from serial dilutions of RNA and were found to be comparable between the two primer pairs. The ^ΔΔ^Ct quantification was performed and the fold change was calculated from 2^−ΔΔCt^. The fold change was normalized to the average fold changes of BSMV treated samples.

#### Phylogenetic analysis of SR proteins

Protein sequences of the cloned *Sr* genes were download from GenBank: SR9B (UYX79454.1), SR13 (ATE88468.1), SR21 (AVK42832.1), SR22 (CUM44200.1), SR22B (UFE16541.1), SR26 (QKW90242.1), SR27 (QZA87369.1), SR33 (AGQ17384.1), SR35 (QNU41031.1), SR45 (QNU41028.1), SR46 (AYV61514.1), SR50 (ALO61074.1), SR61 (QKW90243.1), and SR66 (QXY82434.1). MEGA12^98^ was used to build a phylogenetic tree with MUSCLE alignment and the Maximum Likelihood method (1000 bootstrap replicates).

### Synteny analysis of *SrAect1* locus

Five proteomes of the primary transcripts were downloaded from Phytozome (*B. distachyon* v3.2, *H. vulgare* Morex V3, *O. sativa* v.7.0, *So. bicolor* v5.1, and *Z. mays* B73 NAM-5.0.55); one HC proteome from GrainGenes (*S. cereale* Rye_Lo7_2018_v1p1p1); and three additional proteomes from the original publications (*Ae. sharonensis*^68^, *Ae. tauschii* v6.0^23^, and and *T. aestivum* IWGSC Refseq v2.1^57^). Orthologous and synteny analysis was performed using GENESPACE (v1.3.1)^71^, as described above. Protein domains were identified using InterProScan^66^, as described above. The *SrAect1* locus specific SANT/Myb domain encoding genes are *AE.SHARON.r1.7SG0651220*, *AET7D6G0500000, TraesCS7D03G1275100*, *TraesCS7B03G1286300, TraesCS7A03G1347400*, *SECCE6Rv1G0447180*, *HORVU.MOREX.r3.7HG0751030*, *Bradi1g29680*, *LOC_Os06g51260*, and *Zm00001eb228670*.

## Acknowledgements

We are grateful to Toni Mohr and Roger Thilmony (USDA-ARS, Albany, CA, USA) for help with RT-qPCR; This research used resources from the SCINet project and the AI Center of Excellence of the USDA Agricultural Research Service, under ARS project numbers 0201-88888-003-000D and 0201-88888-002-000D. This work was supported by the ARS CRIS project number 2030-21430-015-000D, and an appointment to the ARS Research Participation Program administered by the Oak Ridge Institute for Science and Education (ORISE) through an interagency agreement between the United States Department of Energy (DOE) and USDA. ORISE is managed by Oak Ridge Associated Universities (ORAU) under DOE contract number DE-SC0014664. Mention of trade names or commercial products in this article is solely for the purpose of providing specific information and does not imply recommendation or endorsement by USDA.

## Author contributions

Y.Q.G., S.S.X., and M.-C.L. initiated and conceived the study. H.-L.Y., N.H., H.-C.C., M.-C.L., S.S.X., and Y.Q.G. planned, organized, planted, collected samples, and conducted the sequencing library preparation and managed sequence data; H.-L.Y., M.-C.L., X.F.Z., Y.Q.G., and S.S.X. planned and carried out BNG optical mapping and analyses; H.-L.Y. and S.S.X. conducted the crosses and established the mapping population. H.-L.Y., H.-C.C., and S.S.X. managed the population growth in the greenhouse. H.-L.Y. and M.-C.L. performed data analysis and linkage map construction. P.C., M.-C.L., and Y.Q.G. planned, conducted, and analyzed gap filling, pseudomolecules, annotations, and assemblies; P.C and Y.Q.G., and J.D. performed genome comparative analysis and evolution. H.-L.Y., P.C., and S.S.X. managed diversity panel and analyzed population diversity; Q.J.Z. and P.C., H.-L.Y., and S.M.Y. conducted chromosome analysis; G.Q.L. and X.Y.X. conducted phenotyping for powdery mildew; D.K., S.G., P.O.F., Y.J., A.P.H., S.B.Z., U.G., and T.L.F. contributed to stem rust and leaf rust phenotyping; F.M. and Z.H.L. conducted phenotyping for bacterial leaf streak, tan spot; S.S. and X.Y.X. performed greenbug phenotyping, K. M. A and A.P.H contributed to Hessian fly phenotyping, E.Y and T.Z.S. contributed to data repository at GrainGenes; P.C., H.-L.Y., S.N.M., J.B., M.J.M., and S.S.X. conducted VIGS; H.-L.Y. and P.C. Y.Q.G., S.S.X., and M.-C.L. drafted the manuscript; All authors read, commented on, and approved the final version of the manuscript.

## Completing interests

The authors declare no completing financial interests.

## Data availability

The raw PacBio reads for genome assembly and the transcriptome data (RNA-seq and Iso-seq) from the 16 tissue types for genome annotation were submitted to NCBI under BioProject number PRJNA1220272. The Aecau_v1 assembly was assigned NCBI Genome accession JBLNHV000000000 with the version number JBLNHV010000000. Access to BLAST and genome browser of Aecau_v1 is available at GrainGenes (https://wheat.pw.usda.gov and https://graingenes.org/jb?data=/ggds/ae-caudata). The Aecau_v1 assembly and annotation files were deposited at Dryad (https://doi.org/10.5061/dryad.2bvq83c3p). The raw PacBio reads and assembly (accession JBREWA000000000; version JBREWA010000000) for line 6-0015 were submitted to NCBI under BioProject number PRJNA1330315. The WGS data of the 95 diversity accessions were submitted to NCBI under BioProject number PRJNA1330229. Seeds of *Ae. caudata* accessions used in this study can be requested by contacting the corresponding author.

---

**Extended Data Fig. 1.**
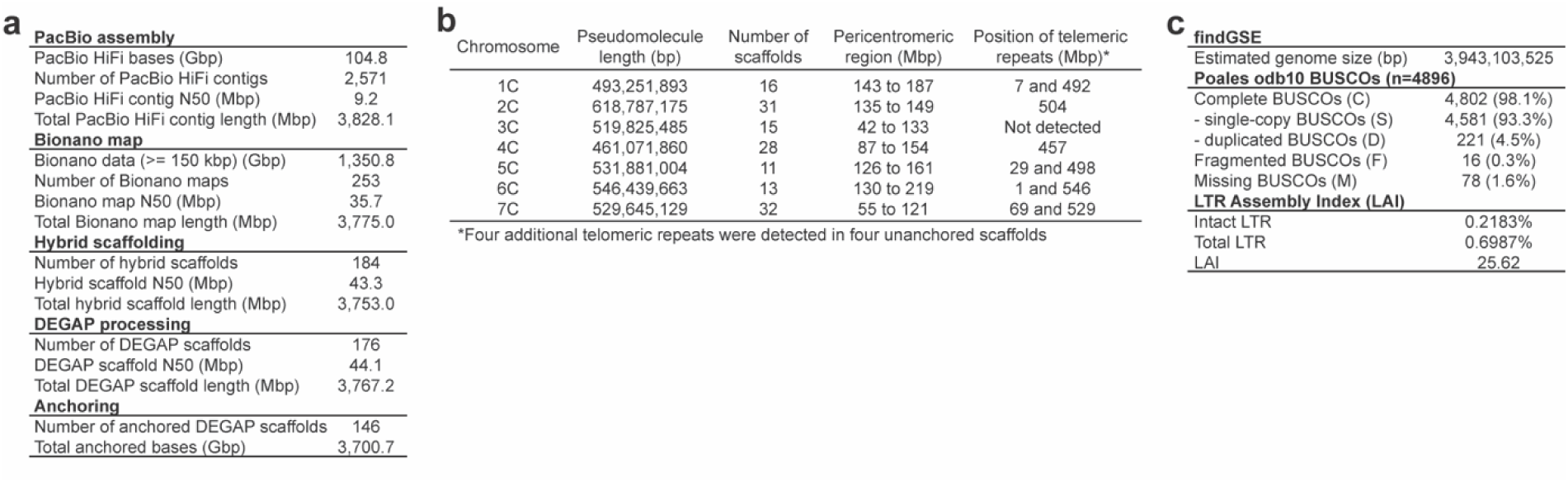
Genome assembly statistics. **a,** Statistics summary of PacBio and Bionano data and assembly at different stages. **b**, Characteristics of the *Ae. caudata* genome Aecau_v1 assembly. **c**, Assembly quality assessments. findGSE estimated the genome size, BUSCO identified conserved orthologs, and LAI reported the intactness index of LTR.

**Extended Data Fig. 2.**
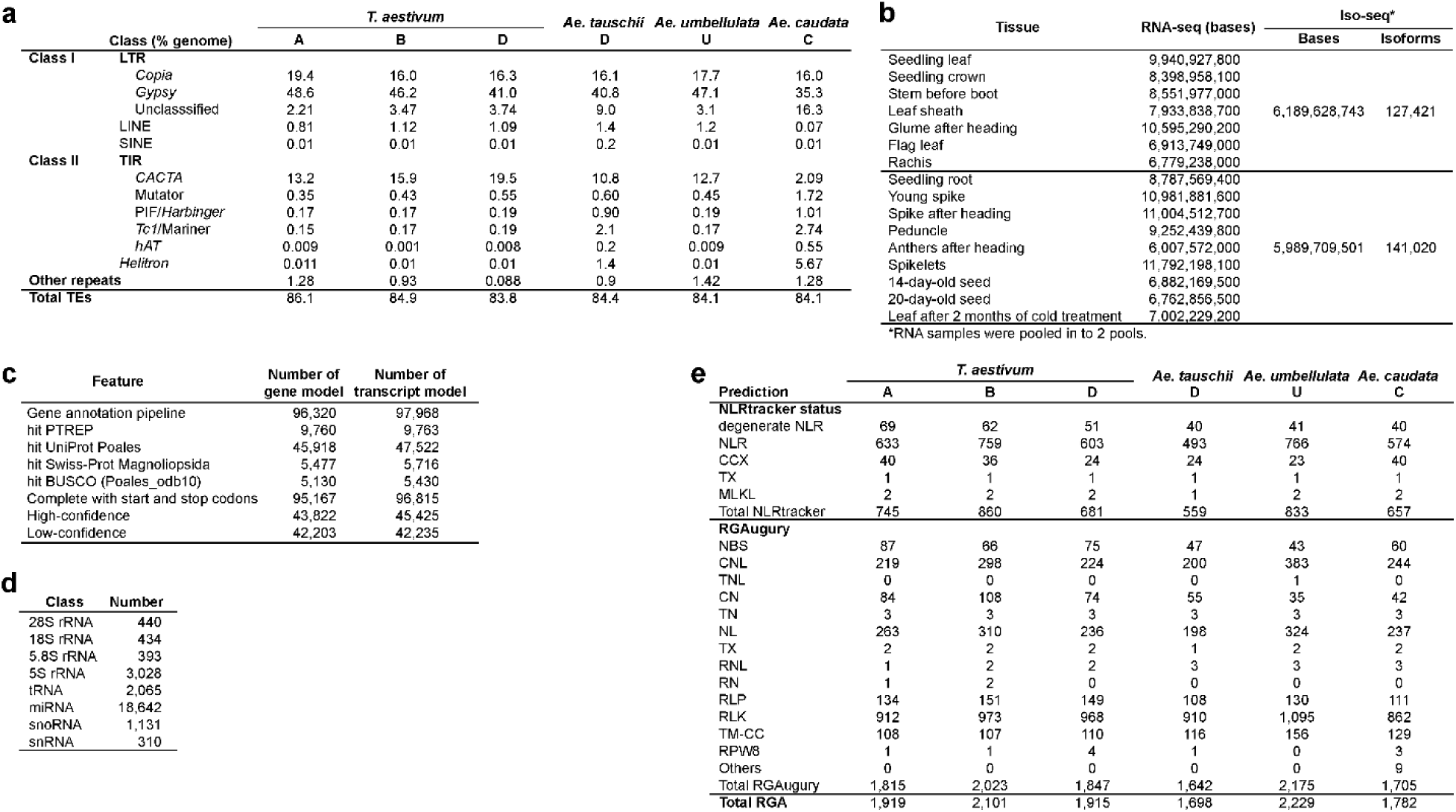
Aecau_v1 annotation statistics. **a,** Transposable element annotation compared to the published results of *T. aestivum* (IWGSC CS RefSeq v2.1), *Ae. tauschii* (Aet v4.0) and *Ae. umbellulata* (PI 554389). **b,** RNA-seq and Iso-seq samples and sequencing summary. **c**, Summary of protein-coding gene annotation and the confidence classification. **d**, Summary of non-coding RNA gene annotation. **e**, Summary of disease resistance gene analog (RGA). Numbers of RGA shown were all derived from the HC gene category. The prediction class abbreviations are based on the NLRtracker and RGAugury programs. NLRtracker classes are NLR (NB-ARC with LRR), degenerate NLR (either RxN-type CC, late blight resistance protein R1, RPW8-type CC, or TIR with a P-loop containing nucleotide hydrolase domain, or a RxN-type CC, late blight resistance protein R1, RPW8-type CC, TIR, or LRR with a RNBS-D, linker, and/or MHD motif), TX, CCX, RPW8 classes are lacking a P-loop containing nucleotide hydrolase domain and RNBS-D, linker, and/or MHD motif, TX (TIR), CCX (RxN-type CC or late blight resistance protein R1), RPW8 (RPW8-type CC), MLKL (HeLo domain of plant-specific mixed-lineage kinase domain like proteins). RGAugury classes are NBS (nucleotide-binding site), CNL (CC-NBS-LRR), TNL (TIR-NBS-LRR), CN (CC-NBS), TN (TIR-NBS), NL (NBS-LRR), TX (TIR-unknown domain), RNL (RPW8-NBS-LRR), RN (RPW8-NBS), RLP (Receptor like protein), RLK (Receptor like kinase), TM-CC (Transmembrane-coiled-coil), RPW8 (Resistance to powdery mildew 8), Others (Other NBS proteins).

**Extended Data Fig. 3.**
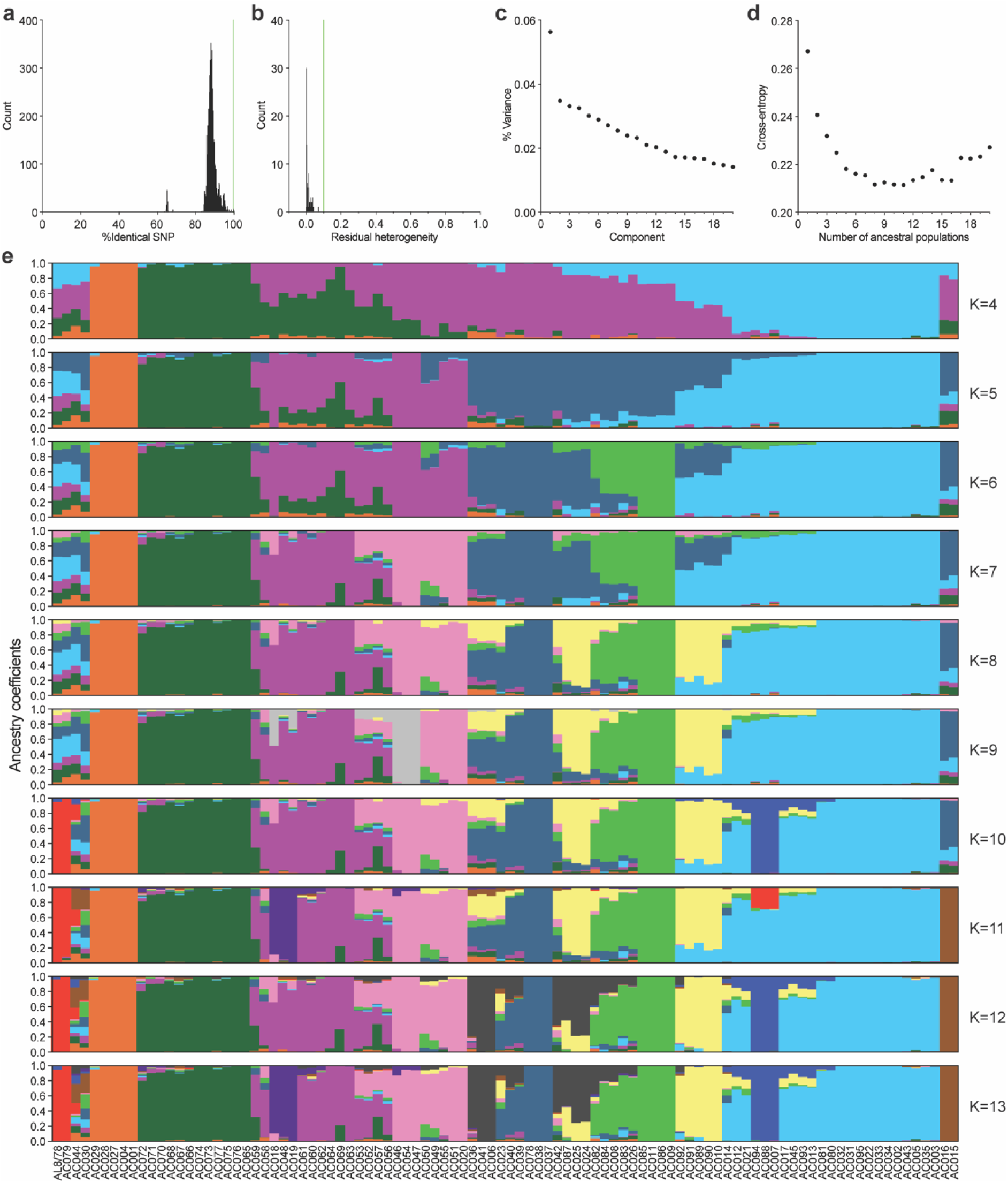
Analysis of diversity panel. **a,** Pairwise comparison of SNP identity percentages to identify redundancy in the panel. The cutoff value of 99.5% (green line) was used. **b**, Residual heterogeneity analysis. All the accessions show values well below the 0.1 threshold (green line). **c**, Variance explained by each component from principal component analysis (PCA). **d**, Cross-entropy analysis. **e**, Ancestry coefficients for K values from 4 to 13.

**Extended Data Fig. 4.**
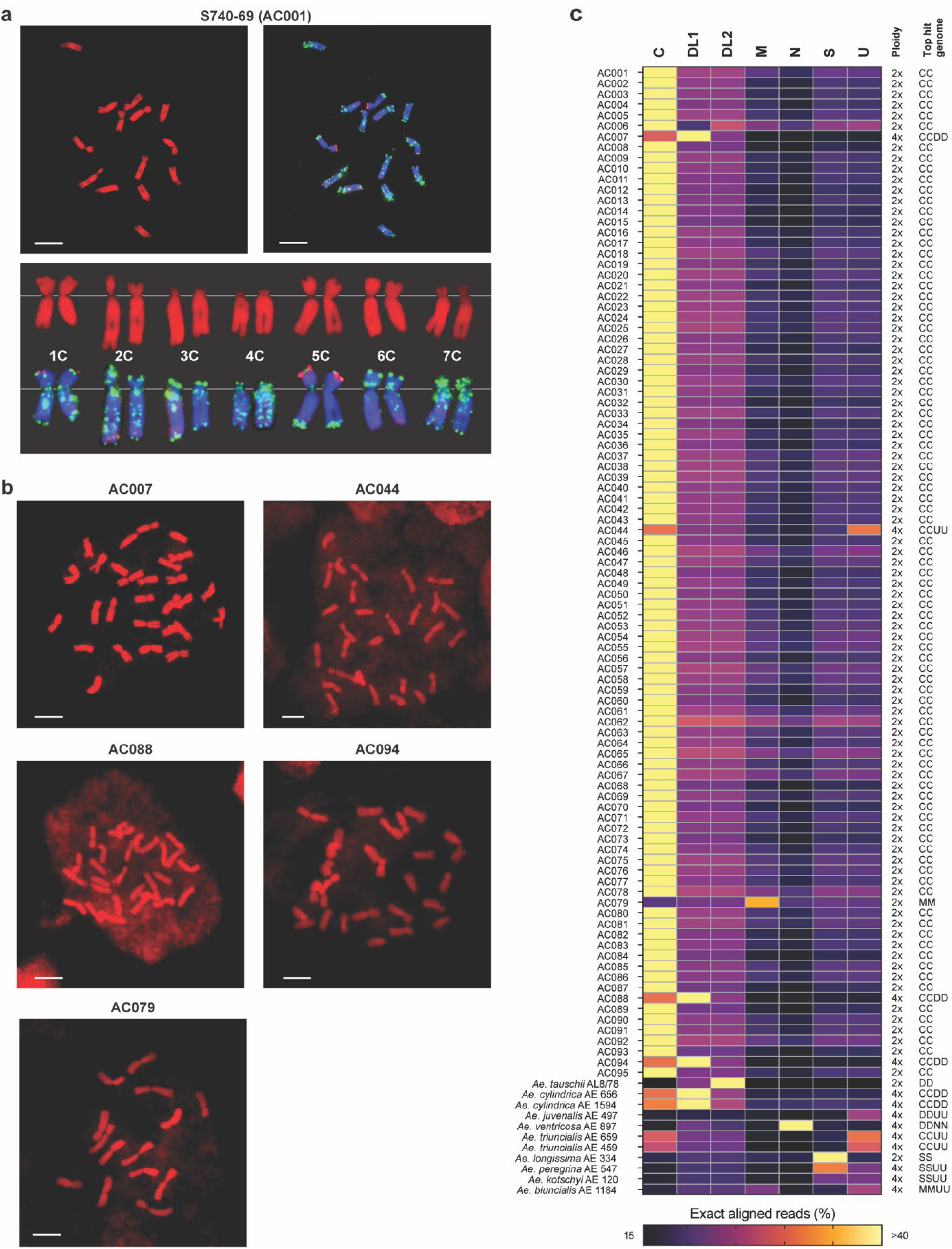
Cytogenetic and sequence analyses of the diversity panel accessions. **a**, *Ae. caudata* S740-69 (AC001) chromosomes. Top left, propidium iodide stained chromosomes. Top right, FISH pattern showing propidium iodide (blue), pSc119.2-1 and GAA probes (green), and pAs1-1, pAs1-3, pAs1-4, pAs1-6, AFA-1, and AFA-4 probes (red). Bottom panel shows the karyotype of S740-69 with chromosome designations. Scale bars = 10 μm. **b**, Chromosomes of tetraploid accessions (AC007, AC044, AC088, and AC094), and diploid accession AC079, which lacks the acrocentric chromosomes observed in *Ae. caudata* C chromosomes (AC001). Scale bars = 10 μm. **c**, Percentage of exactly matched WGS reads from each accession aligned to different *Aegilops* genomes including C (accession S740-69), D (L1) (Lineage 1; TA10171), D (L2) (Lineage 2; AL8/78), M (PI 551049), N (only the N genome of RM271), S (AEG-6782-2), and U (TA1851).

**Extended Data Fig. 5.**
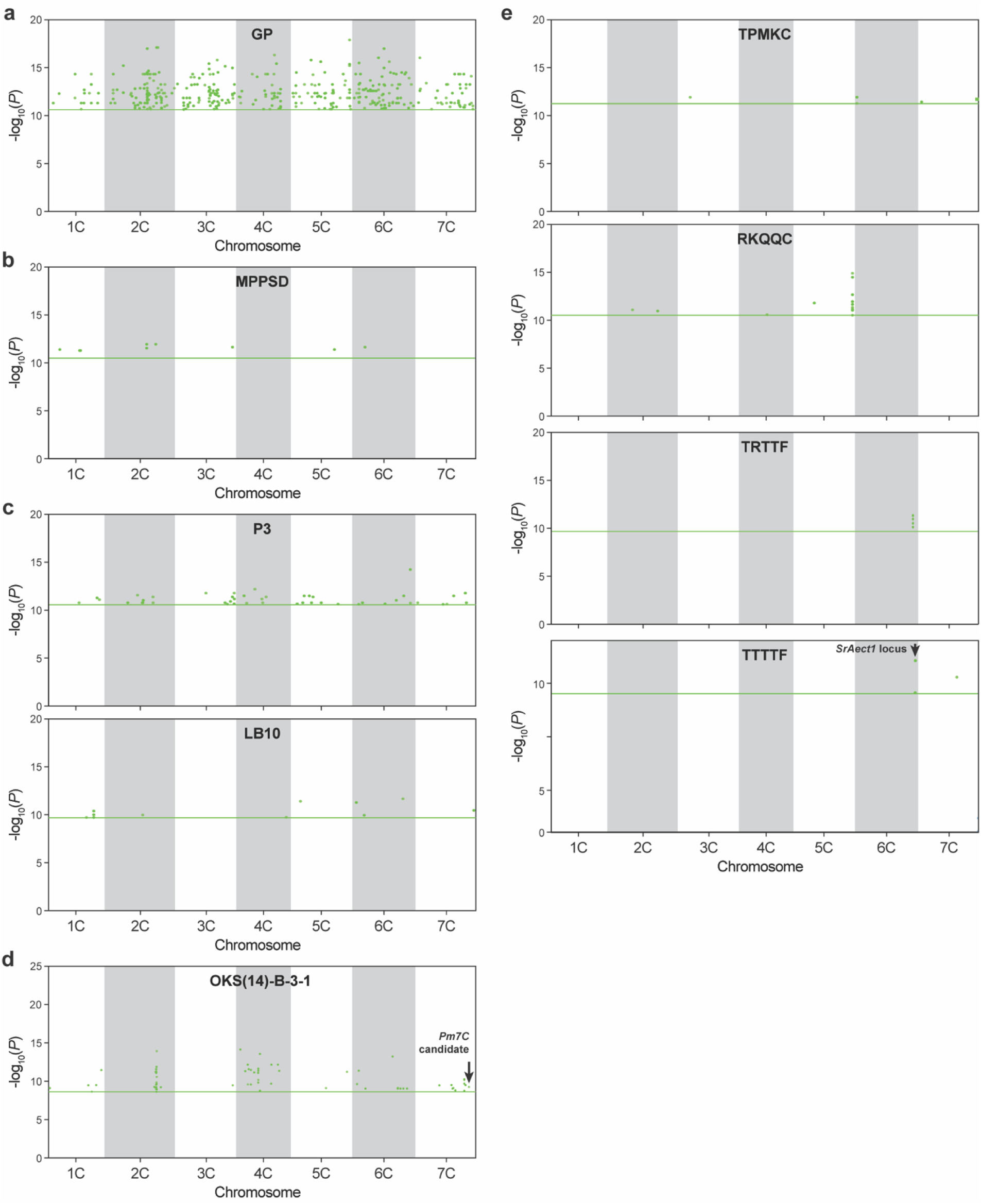
GWAS analysis of insect pest and disease resistance. **a,** Hessian fly (*Mayetiola destructor* biotype GP). GWAS analysis produced >6,000 significant k-mers but <2,000 k-mers were mapped to the reference genome across all chromosomes. Note that the reference genome derived from a susceptible accession. **b,** Leaf rust (*Puccinia triticina* race MPPSD). **c,** Bacterial leaf streak (*Xanthomonas translucens* pv. *undulosa* isolates P3 and LB10). **d,** Powdery mildew (*Blumeria graminis* f. sp. *tritici* isolate OKS(14)-B-3-1). A major peak on chromosome 2C. Some significant k-mers clusters were mapped to the powdery mildew resistance locus, *Pm7C*. **e,** Stem rust (*Puccinia. graminis* f. sp. *tritici* races TPMKC, RKQQC, TRTTF and TTTTF). TTTTF identified significant k-mers within the *SrAect1* locus on chromosome 6C similar to the TTKSK 6C peak. TRTTF identified a different peak on chromosome 6C. RKQQC identified a peak on chromosome 5C. Horizontal lines show the thresholds for the 5% family-wise error rate.

**Extended Data Fig. 6.**
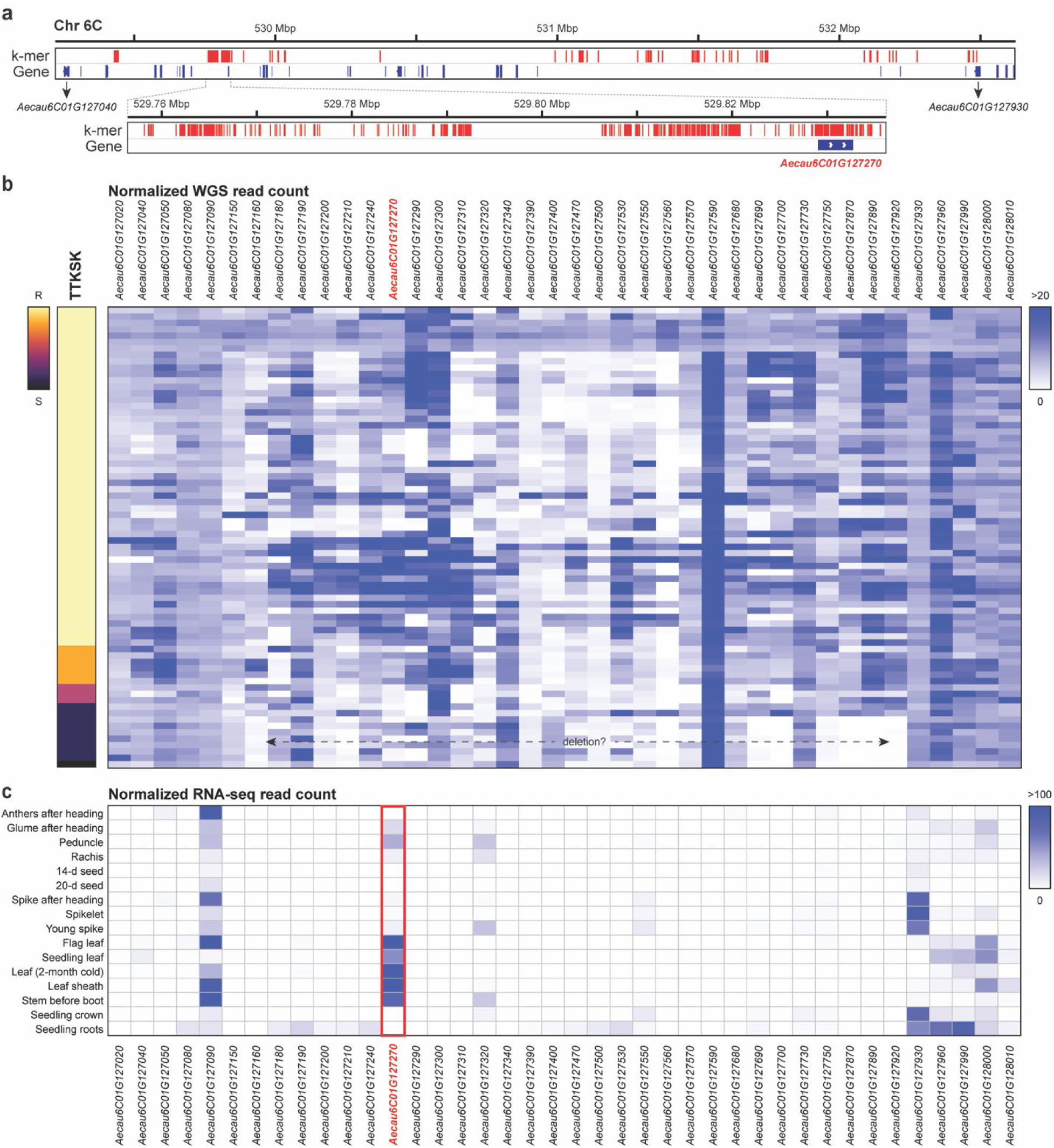
Evidence supporting *Aecau6C01G127270* candidacy. **a**, Distribution of the significant k-mers in TTKSK GWAS analysis within the peak region of the *SrAect1* locus. *Aecau6C01G127270* contains the largest number of significant k-mers. **b**, WGS coverage of the genes in the *SrAect1* locus. TPM normalized counts are shown as a blue-white heatmap. Each row represents an accession. Accessions were grouped based on TTKSK resistance phenotype (leaf heatmap). Accessions that are susceptible to TTKSK show low read coverage across ∼29 genes, suggesting the possibility of their deletions. **c**, Expression patterns of the genes within the *SrAect1* locus. *Aecau6C01G127270* is the only gene in the putative deletion region with strong expression at sites of stem rust infection (leaf, leaf sheath, stem). TPM normalized counts are shown.

## Supplementary Tables

Supplementary Table 1. *Aegilops caudata* diversity panel accession passport details, redundant analysis, chromosome counting, and their resistance to greenbug, Hessian fly, tan spot, leaf rust, bacterial leaf streak (BLS), powdery mildew, and stem rust.

Supplementary Table 2. Linkage map used for scaffold anchoring.

Supplementary Table 3. Assembly statistics and chromosome anchoring coordinates.

Supplementary Table 4. List of high-confidence genes.

Supplementary Table 5. InterPro and Pfam annotations of the high-confidence genes.

Supplementary Table 6. Predicted disease resistance gene analogs.

Supplementary Table 7. Residual heterogeneity.

Supplementary Table 8. Percentage of identical SNP among the accessions

Supplementary Table 9. Ancestry coefficient analysis.

